# Defining amino acid pairs as structural units suggests mutation sensitivity to adjacent residues

**DOI:** 10.1101/2022.10.23.513383

**Authors:** Aviv A. Rosenberg, Nitsan Yehishalom, Ailie Marx, Alex Bronstein

## Abstract

Proteins fold from chains of amino acids, forming secondary structures, α-helices and β-strands, that, at least for globular proteins, subsequently fold into a three-dimensional structure. A large-scale analysis of high-resolution protein structures suggests that amino acid pairs constitute another layer of ordered structure, more local than these conventionally defined secondary structures. We develop a cross-peptide-bond Ramachandran plot that captures the conformational preferences of the amino acid pairs and show that the effect of a particular mutation on the stability of a protein depends in a predictable manner on the adjacent amino acid context.

**One-Sentence Summary:** Large-scale protein backbone analysis reveals amino acid pair conformational preferences and predicts how sequence context affects mutant stability.

---

Protein function is dictated by structure, which can be globular – rigid, ordered and compact, or intrinsically disordered. Either way, proteins must adopt conformation(s) in a predictable manner supporting reproducible interactions in the cellular network. This suggests that random coil regions of the amino acid chain may not be so random and points towards the existence of order at a level more local than the well-defined secondary structures, helices, strands and turns. Evidence for such local structure include a work by Fitzkee and Rose (2004) who presented what they termed a “physically absurd model” showing that varying the backbone dihedral angles in only 8%of the residues in structured proteins created proteins with random coil properties, in terms of end-to-end distances and mean radii of gyration (1). An extensive investigation carried out by Eker et al (2004) showed that alanine-based tripeptides, AXA, have intrinsic structural propensities dependent on the identity of amino acid X (2). More recently, structural preferences and resilience have been shown on disordered proteins in vitro and in cell (3). The first attempts to capture protein backbone structure beyond the well-recognized secondary structures were made more than thirty years ago. Unger et al (1989) showed that six amino acid-long sections of the backbone cluster into about 100 specific conformations and that entire protein structures can be well reconstructed from these building blocks (4). The idea of creating such structural alphabets from the common conformations formed by consecutive residues has since been explored and used extensively, commonly as “Protein Blocks” of 16 prototypes defined by five consecutive amino acids (5–7). Alternative to the fragment-based approach, attempts to capture local structure have included describing rotation, twist and rise-per-residue parameters in helical structures (8), using tripeptides and a measurement of their rigidity (9), and defining a (φ, ψ)_2_ motif (10–11). The (φ, ψ)_2_ motif defines amino acid pairs as a structural elements and is attractive in being very local and restricted to a discrete set of 400 elements. We suggest a different angle, literally, on defining amino acid pairs as structural elements and analyzing their preferred backbone conformations. We show that amino acid pair conformations are well captured in an “amino-domino” model; each pair is described by only two dihedral angles, those separated by the peptide bond, with each amino acid contributing to two dominos, in one domino being described by ψ and in another by φ. We denote these cross-bond angle pairs as (ψ_*k*_, φ_*k*+1_) where k is an index of an amino acid. Two important advantages of the amino-domino model are that (a) all atoms defining (ψ_*k*_, φ_*k*+1_) are part of the amino acid pair, making it the smallest self-contained structural unit; and (b) the description of the amino acid pair remains two-dimensional and has a clear geometric interpretation that can be captured using the familiar Ramachandran plot, but of these two angles (ψ*_k_*, φ_*k*+1_). Exploring the biological implication of this model, we demonstrate that single point mutations are sensitive to the identity of the adjacent residues in the amino acid chain in a manner which is predictable by comparison of the (ψ*_k_*, φ_*k*+1_) distributions of the relevant amino acid pairs.

The protein backbone is commonly described as a series of dihedral angle pairs, (φ*_k_*, ψ*_k_*). The joint distributions of these angle pairs fall into specific (allowed) regions, compatible with the stereochemical constraints of the molecule, and corresponding to the backbone conformations which are characteristic of the protein secondary structures as shown by Ramachandran almost sixty years ago (11). It is of note that although (φ*_k_*, ψ*_k_*) are conventionally assigned to an individual amino acid, they are not entirely defined by that amino acid, since each of the dihedrals relies on the coordinates of an atom from each of the adjacent residues. Seeking to identify the smallest, *self-contained* structural element possible we considered describing the protein backbone as the series of overlapping amino acid pairs, analogous to dominos, and captured by the dihedral angle pairs (ψ*_k_*, φ_*k*+1_). It is of note that such pairs of angles are separated by the peptide bond and so have less overlap of contributing atoms. Specifically, the two adjacent dihedrals, (φ_*k*_, ψ*_k_*), share three of the four atoms which define them, while the pair (ψ*_k_*, φ_k+1_) share only two atoms (Fig. 1, top). Using high-resolution protein structures from the Protein Data Bank (PDB), we calculated the (*ψ_k_*,φ_*k*+1_) dihedral angle distribution and compared it to the traditional (φ_*k*_, ψ_*k*_) dihedral angle distributions (Fig. 1, middle). A clearly observable difference is that the cross-peptide-bond Ramachandran plot of (ψ_*k*_, φ_*k*+1_) covers more of the conformational space than the traditional (φ_*k*_, ψ*k*) plot. More importantly, the variety of local structural conformations that pairs of amino acids assume (Fig. 1, bottom) map into distinctly separate locations in the cross-bond Ramachandran plot (Fig.1, middle-center), whereas the same conformations have overlapping locations in the traditional Ramachandran plot (Fig. 1, middle-left).

**Fig. 1.**
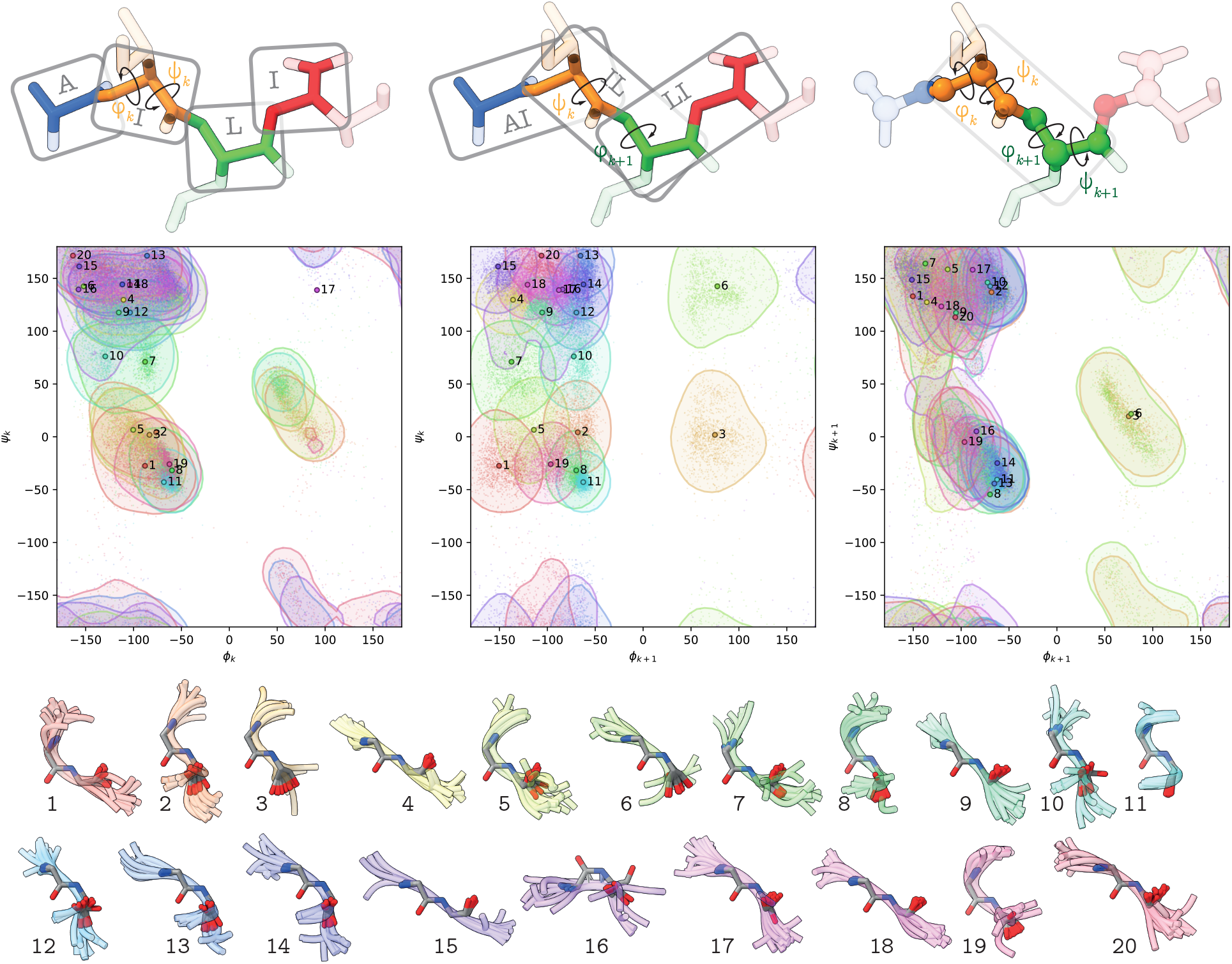
Describing the protein backbone through a different angle pair better explains conformations adopted by amino acid pairs. *Top*: rather than treating each amino acid as a single structural unit described by (φ_*k*_, ψ_*k*_) (top-left), we propose overlapping pairs of amino acids described by (ψ_*k*_, φ_*k*+1_) as a form of structural “domino blocks” (top-center). Traditionally, dihedral angle pairs are defined by the two atoms on either side of the bond around which they are centered, making the dihedral angles (φ_*k*_, ψ_*k*_) describing a single amino acid dependent on atoms from the neighboring amino acids: φ_*k*_ depends on the preceding nitrogen atom N_*k*-1_, while ψ_*k*_ depends on the following carbon atom *C*_*k*+1_ (top-right). Contrastingly, the pair of dihedral angles (ψ*_k_*, φ_*k*+1_) bridged by the peptide bond connecting two consecutive amino acids *k* and *k*+1 depends only on the atoms contained within the pair. *Middle*: pairs of amino acids from 24,875 different proteins are visualized in the form of dihedral angle distributions known as the Ramachandran plot, and assigned to one of 20 clusters. Left-to-right are shown the different two-dimensional marginals of the four-dimensional joint distribution (φ_*k*_, ψ_*k*_, φ_*k*+1_, ψ_*k*+1_), namely, (φ_*k*_, ψ*k*), (ψ*k*, φ_*k*+1_), and (φ_*k*+1_, ψ_*k*+1_). The variety of conformations assumed by the amino acid pairs is clearly discernible in the cross-bond Ramachandran plot (middle-center), unlike the traditional plots (middle-left and right) in which distinct conformations are indistinguishable. Angles are indicated in degrees and have a periodic boundary at ±180°. Contours indicate 90% probability regions. Colors indicate cluster assignment and numbered points indicate their representatives. *Bottom*: A visualization of the backbones from 10 representative amino acid pairs, within each of the 20 clusters depicted in the Ramachandran plots. Context of two surrounding amino acids also shown. Backbones were clustered according to their Euclidean (rigid body) similarity after canonical alignment of atom locations.

The increased occupancy of the two-dimensional conformational space might be expected due to the reduced overlap of defining atoms for angles in the cross-bond pair. Interestingly, transitions between secondary structures can only be discerned by the new plot. A prime example is Cluster 6 which occupies a distinct position in the first quadrant of the cross-bond Ramachandran plot which is not populated in the traditional Ramachandran Plot. This position captures the recognizable and characteristic (ψ_*k*+1_, φ_*k*+2_) conformation of the apex of a type II β-turn which is formed when the *k*+1 and *k*+2 amino acids adopt ψ ~ 120° and φ ~ 80° angles, respectively (12). Through the lens of the cross-bond-Ramachandran it can now be appreciated that this conformation, whilst necessary to form a type II β-turn, it is not specific to this structural context. Analysis of the representative amino acid pair backbones from Cluster 6 reveals that whilst some of the amino acid pairs are indeed located within such well-defined turns, there is a range of other structural contexts from which these pairs are carved (Fig. 2).

**Fig. 2.**
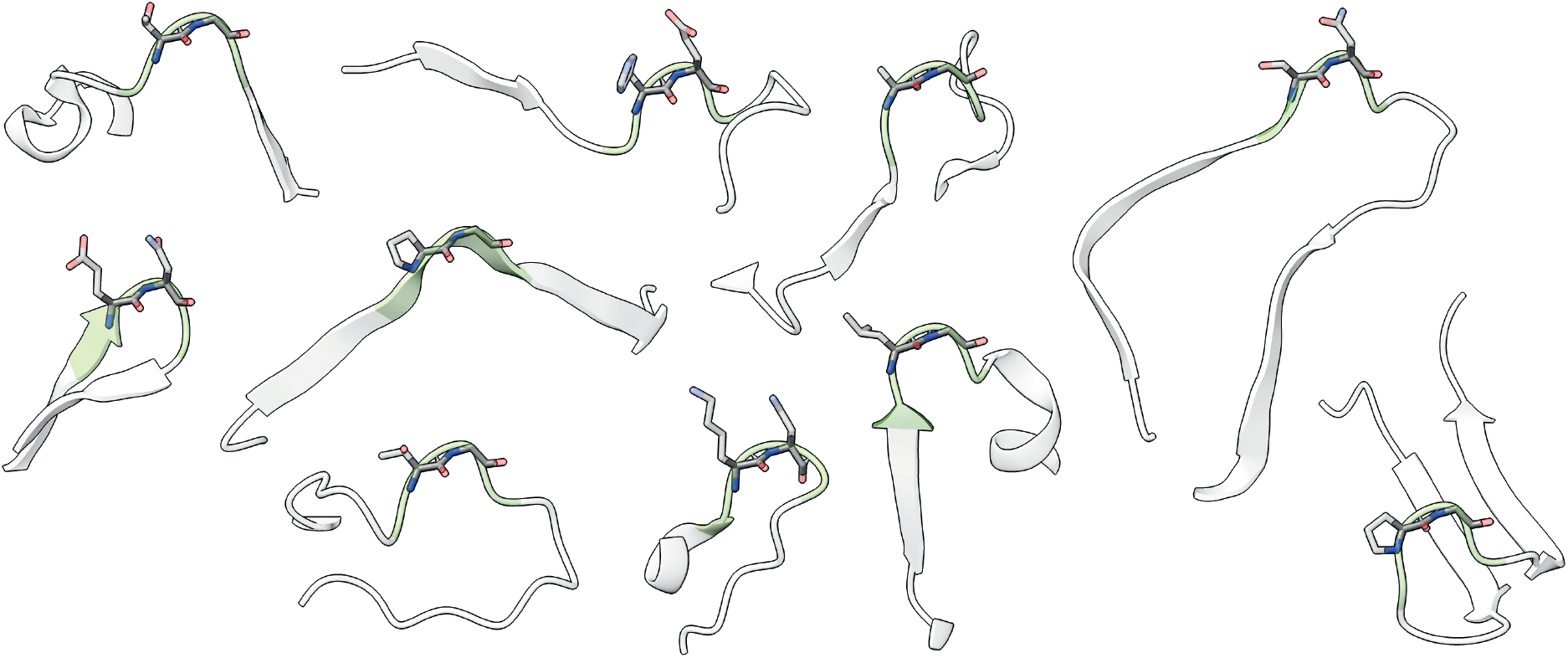
Clusters of amino acid pair conformations encompass a range of larger structural contexts. Amino acid pairs (displayed as sticks) from the ten representative structures of Cluster 6 are shown in their greater structural context (displayed as ribbons).

The larger conformational space occupied by the cross-bond angle pair, (ψ*_k_*, φ_*k*+1_), also indicates that their constituent angles are less statistically dependent, as can be readily explained by the reduced overlap of atoms defining them. Importantly, this greater independence does not imply that the trajectories of cross-bond angles under thermodynamic fluctuations would be less dependent than the trajectories of (φ_*k*_, ψ_*k*_). We performed molecular dynamics simulations of small proteins harboring loops composed of residues not influenced by interactions with the rest of the protein. The results indicate that that the constituents of the cross-bond pair (ψ*_k_*, φ_*k*+1_) are at least as correlated as those of the regular pair (φ_*k*_,ψ*_k_*) (Fig. 3 and Table S1). This can be explained by the rigidity of the peptide bond. When observing a specific cross-bond angle pair (i.e., the distribution is conditioned on specific location in a structure), the rigid peptide bond ensures these two angles produce a highly correlated distribution. In contrast, when observing the cross-bond angle pairs unconditioned on specific locations within structures, lower correlation is observed in the cross-bond distribution compared to the traditional one (Fig. 1 middle-left, center).

**Fig. 3.**
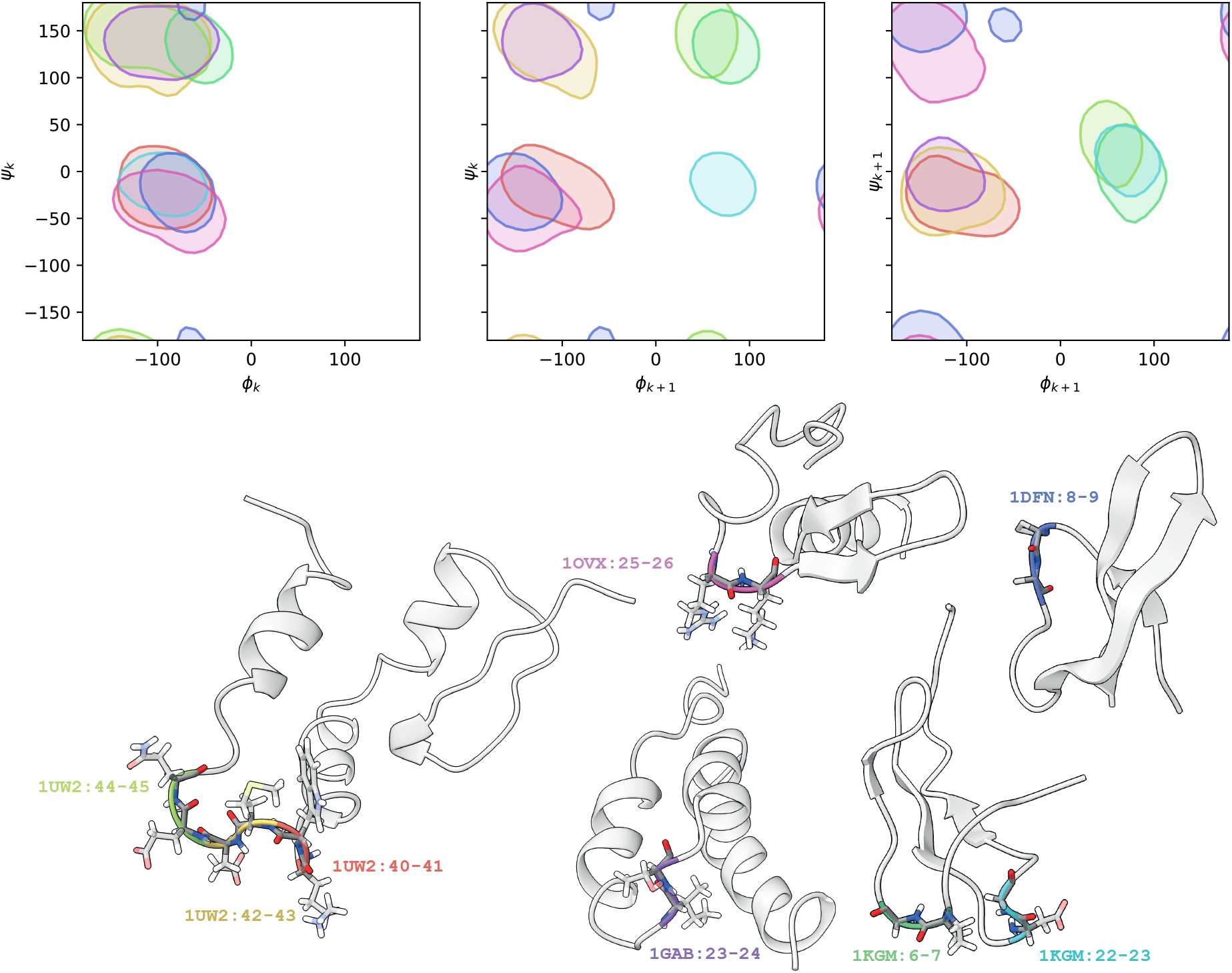
Molecular dynamics simulations demonstrate that given a specific location, the cross-bond dihedral angle pair are as correlated as the traditional dihedral angle pair. *Top* left-to-right: joint distributions of the dihedral angles (φ_*k*_, ψ_*k*_), (ψ_*k*_, φ_*k*+1_), and (φ_*k*+1_, ψ_*k*+1_) of eight non-interacting amino acid pairs in five different protein structures. Contours indicate 90% probability regions. *Bottom*: PDB IDs and residue numbers highlight the locations of the simulated amino acid pairs in the corresponding structures with matching color codes.

An important feature obtained by shifting the conventional reference frame, of the immediately adjacent angle pair, to the cross-bond dihedral angle pair is that the angles are now entirely contained within the amino acid pair it is being used to describe. This allows for context-free assignment of amino acid pair propensities to different conformations or regions on the cross-bond plot. Focusing again on Cluster 6 it is evident that the propensities for the first and second position of this pair (Sup. Fig. 2) correspond with the well characterized propensities for the *k*+2 and *k*+3 positions in β-turns; namely, the first position being most favored by proline and the second position being disfavored by most amino acids except glycine, asparagine and aspartic acid (13). Cluster 11 also exemplifies a clear agreement between the new amino acid pair cluster propensities and known structural conformations. This cluster corresponds to the structure adopted in the middle of an alpha helix, as all three marginals, (φ_*k*_, ψ*_k_*), (ψ*_k_*, φ_*k*+1_), and (φ_*k*+1_, ψ_*k*+1_) of the amino acid pair, have the same mode, which is characteristic of α-helices (Sup. Fig. 2). In accordance, the propensities of this cluster appear similar in the first and second positions of the pair and correlate well with the known amino acid secondary structure propensities (14) for α-helices; higher for alanine, arginine, glutamic acid, leucine, methionine and glutamine and lower for proline, glycine, asparagine, threonine and serine. Most of the other clusters represent specific regions within a secondary structure or transitions between secondary structures. The new perspective proposed here should therefore refine the analysis of studies dedicated to exploring individual amino acid preferences towards different structural elements.

Another implication of the amino-domino model is in demonstrating and quantifying how immediately adjacent amino acids in the sequence can influence the effect of a single point mutation. A mutation alters the side chain at a particular position in the structure, affecting the interaction network and chemistry around that position. Nevertheless, most approaches and tools consider mutations independent of the immediate sequence context, taking mainly the side chain identity into account. At the level of a single amino acid, we find that the distance between the (φ_*k*_, ψ*_k_*) dihedral angle distributions of two amino acids is predictive of their BLOSUM62 score (Fig. S4), possibly capturing how amino acids having similar side chains can be accommodated in similar structural locations. We suggest that beyond the side chain, an additional effect of mutation may result from altered backbone conformational preferences. To probe for how mutations might affect backbone conformational preferences, we calculated the distances between the cross-bond Ramachandran plots of pairs of amino acids and observed that the same amino acid substitution may have vastly different impact on the (ψ*_k_*, φ_*k*+1_) distribution depending on the partner amino acid in the left or right position in the pair. For example, as visualized in Fig. S3, FI and YI have indistinguishable cross-bond distributions, while FH and YH exhibit very distinct distributions. We hypothesized that this distance may be predictive of how structurally disruptive a mutation may be, even in cases where the F>Y side chain substitution has little impact on the environment.

To corroborate this hypothesis, we mutated surface-exposed, non-interacting amino acids in the β- barrel of the green fluorescent protein (GFP). Each mutation X>Y was made in two different adjacent amino acid contexts, A_1_XB_1_>A_1_YB_1_ and A_2_XB_2_>A_2_YB_2_. For the first context, A_1_ and B_1_ were selected to minimize the distance between the cross-bond Ramachandran plots of (A_1_X, A_1_Y) and (XB_1_, YB_1_), while for the second context A_2_ and B_2_ were selected to maximize the distance between (A_2_X, A_2_Y) and (XB_2_, YB_2_). Melting temperature ΔT_m_ relative to the wild type was used to quantify the change in protein stability. Fig. 3 shows that the distance between cross-bond Ramachandran plots is predictive of ΔT_m_.

**Fig. 3.**
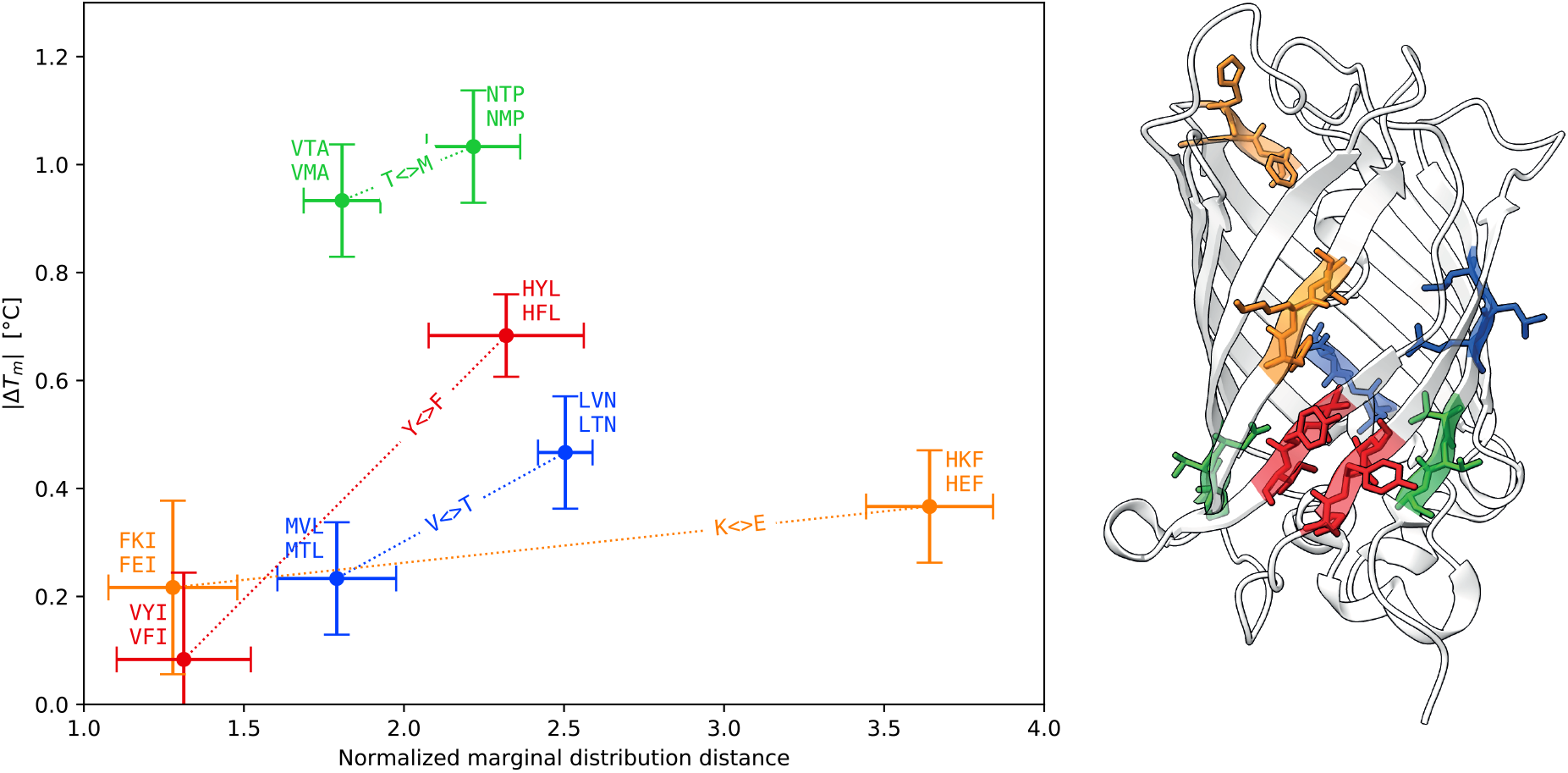
Similarity of cross-bond Ramachandran plot is predictive of the impact of a conservative mutation on protein stability. Four conservative mutations were made in the GFP (PDB:2B3P) in contexts either minimizing or maximizing distances between (ψ_*k*_, φ_*k*+1_) dihedral angle distributions of two consecutive amino acid pairs containing the mutation. For example, in VYI>VFI, VY>VF and YI>FI have similar cross-bond Ramachandran plots, while in HYL>HFL, HY>HF and YL>FL have dissimilar plots. Protein stability, quantified as ΔT_m_ relative to the wild type, is correlated to the distance between the cross-bond angle distributions. Distance is defined as the maximum over the normalized distance between the left and the right pairs including the mutation (e.g., for VYI>VFI, it is the maximum over the distances between VY, VF and YI, FI). Confidence intervals of 1σ are shown.

Defining amino acid pairs as structural units that are effectively described by the unconventional, across-the-peptide bond pair of dihedral angles provides a useful and readily accessible tool for probing and predicting local structural preferences in both native and mutant proteins and may find particular use in structural studies on disordered proteins and regions or in homology modeling.

## Author contributions

AR, AB and AM conceived the idea, designed the methodology and interpreted the results. AR and AB conducted the data analysis. NY and AM conducted the GFP experiments.

## Data and materials availability

Data and code will be made available before publication

## Supplementary Materials

### Materials and Methods

Most of the analysis procedure is based on the data collection and mathematical tools described in the Methods section of Rosenberg et al. (15). Here we highlight mainly the additional details and modifications.

#### Protein structures collection

Non-redundant protein chains were collected using the method described in (13), with a slightly different initial PDB query. The query parameters were (i) Method: X-Ray Diffraction; (ii) X-Ray Resolution: Less than or equal to 1.8 Å; (iii) R_free_: Less than or equal to 0.24, (iv) Expression system contains the phrase “*Escherichia Coli*”. The PDB structures were processed using the biopython package in order to obtain the backbone-atom coordinates at each residue. In total, we collected 5,648,759 consecutive amino acid pairs from 24,875 distinct PDB protein chains, discarding all pairs that were closer than 5 positions to the chain termini.

#### Clustering of amino acid pairs

In order to obtain the 20 clusters depicted in figs 1 and S2, each amino acid pair was first brought into a canonical orientation (Fig. S1). This was achieved by setting the location of the C atom of the first amino acid at the origin, aligning the peptide bond in the direction of the *x* axis, and the plane formed by the C^α^ and C atoms of the first amino acid and the N atom of the second amino acid with the *xy* plane, with the normal pointing in the *z* axis direction. The coordinates of the canonically oriented C^α^, C, and N atoms of both amino acids were then stacked into an 18-dimensional vector representing the pair. The set of all such vectors from all the pairs was then clustered into *k*=20 clusters using Lloyd’s *k*-means algorithm with the Euclidean metric implemented in the `sklearn` python package. 10 runs with different centroid seed initializations were used, with the maximum of 300 iterations in each run. Amino acid pairs closest (in the Euclidean sense) to the calculated cluster centers were selected to represent the cluster. Each cluster was further clustered into 10 sub-clusters using the same procedure to obtain the 10 representative backbones depicted in Fig. S2. Although the ordering of the clusters is unimportant, we used a multidimensional scaling (MDS) procedure to ensure that closely-indexed cluster centers are closer to each other in the sense of the above-defined Euclidean distance.

**Fig. S1.**
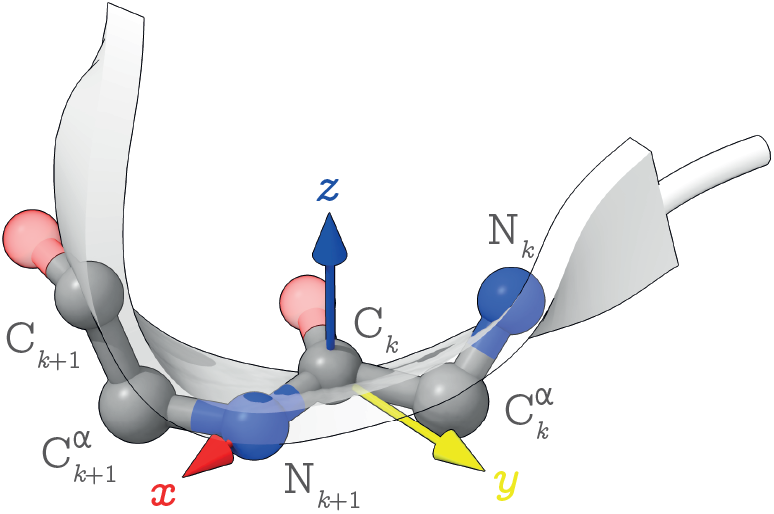
Canonical alignment of amino acid pair conformations. The C atom of the first amino acid is brought to the origin, the peptide bond to the consecutive amino acid is aligned with the *x* axis, and the normal to the C^α^, C, N’ plane is aligned with the positive direction of the *z* axis.

#### Calculation of cluster propensities

The propensity *π*(A * |*c*) of an amino acid pair `A*` (with the star indicating that any amino acid can occupy the second position) to cluster *c* (reported in Fig. S2) was calculated as the log of the ratio of the probabilities which were, in turn, estimated from frequencies:

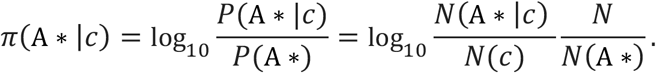

Here *N* denotes the total number of pairs, *N*(*c*) the size of cluster *c*, *N*(*A* *) the total number of pairs with `A` in the first position, and *N*(*A*\*|*c*) the number of pairs with `A` in the first position in cluster *c*. The propensities of amino acid pairs of the form `*A` were calculated in the same manner.

#### Dihedral angle density estimation

Dihedral angle densities depicted in the Ramachandran plots (figs. 1 and 3) were calculated using the kernel density estimation (KDE) procedure described in detail in (13). We used square bins of width 2° and a Gaussian kernel with σ=12°. Regions containing 90% probability and contours containing 25%, 50%, and 75% of the probability (Fig. S3) were calculated using the procedures detailed in (13).

#### Calculation of distances between Ramachandran plots

Distances between Ramachandran plots were estimated as L_1_ distances between KDEs estimated from their points. To gauge the effect of finite sample sizes, average distances and their standard deviations were calculated on 100 random samples of 1000 angle pairs bootstrapped independently with replacements.

To calculate the normalized distances between pairs of amino acids `AX` and `AY` reported in figs 3 and S4, we calculated the ratio of the L_1_ distances between the corresponding KDEs and the KDEs of `*X` and `*Y`,

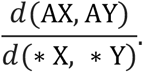

Where the KDEs of `*X` and `*Y` are estimated from points of any pairs with `X` and `Y` at the second position, respectively. As with the unnormalized distances, 100 random samples of 1000 angle pairs bootstrapped independently with replacements were used to estimate the confidence intervals.

#### Calculation of correlation coefficients between dihedral angles

Correlation coefficients reported in Table S1 were calculated on the MD simulation data from Fig. 3 using the flat torus statistics described in detail in (13).

#### Multidimensional scaling (MDS) plots

The similarity plots in Fig. S3 were calculated using the probabilistic MDS procedure proposed and described in (13).

#### Molecular dynamics simulations

Proteins having a short sequence length were randomly selected and inspected in pymol for the presence of loops regions having at least three amino acids in sequence making no polar interactions with the rest of the protein. Single chains without water molecules were prepared for molecular dynamic simulations using the PDB2PQE server (https://server.poissonboltzmann.org/, 16). Molecular dynamics was preformed using GROMACS (17,18). Prior to measuring dynamics an equilibrated and energy minimized system was prepared by standard procedures including solvation and ion addition to achieve a neutral state. NVT (constant Number of particles, Volume, and Temperature) was carried out for 100-ps and 300K. NPT (constant Number of particles, Pressure, and Temperature) was carried out for 100-ps. We ran 10-ns MD simulations with trajectory images being recorded every 10-ps. Relevant backbone φ and ψ angles were extracted and analysed.

#### Wild type and mutant GFP melting temperature measurments

sfGFP his-tagged at both the N and C terminals in expression plasmid pET28a was obtained as a gift from Avi Schroeder. Point mutations were introduced using complementary primers designed in the NEBaseChanger program and following the standard Q5 site directed mutagenesis protocol. The introduction of mutations were verified by sequencing. Wild type and mutant (V120T, V219T, K26E, K166E, T186M, T225M, Y151F and Y200F) sfGFP protein was overexpressed in BL21 cells. A 10mL overnight starter culture was added to 500mL LB supplemented with kanamycin and grown to OD600 ~ 0.6. Protein expression was induced by the addition of 1mM IPTG and cells were grown for a further 4 hours before being collected by centrifugation and stored at −20°C until purification. Frozen cells were resuspended in lysis buffer (150mM NaCl, 50mM Tris pH 8) before disruption in a microfluidizer. Following centrifugation to remove cellular debris the supernatant was passed through a nickel column which was then first washed with 10CV lysis buffer and then another 5CV lysis buffer supplemented with 10mM imidazole. The protein was eluted in lysis buffer containing 250mM imidazole which was subsequently removed by overnight dialysis against lysis buffer. The purity of the protein was confirmed by SDS-PAGE and the protein samples were concentrated to 5mg/ml. Melting temperature was determined by differential scanning fluorimetry, measuring the fluorescence every 0.5°C upon heating from 20°C to 95°C using 80% power in a Prometheus NT.48 nano-DSF instrument (Nanotemper Technologies). The results shown in Fig. 3 represent 5 independent measurements from each of two biological repeats.

**Fig. S2.**
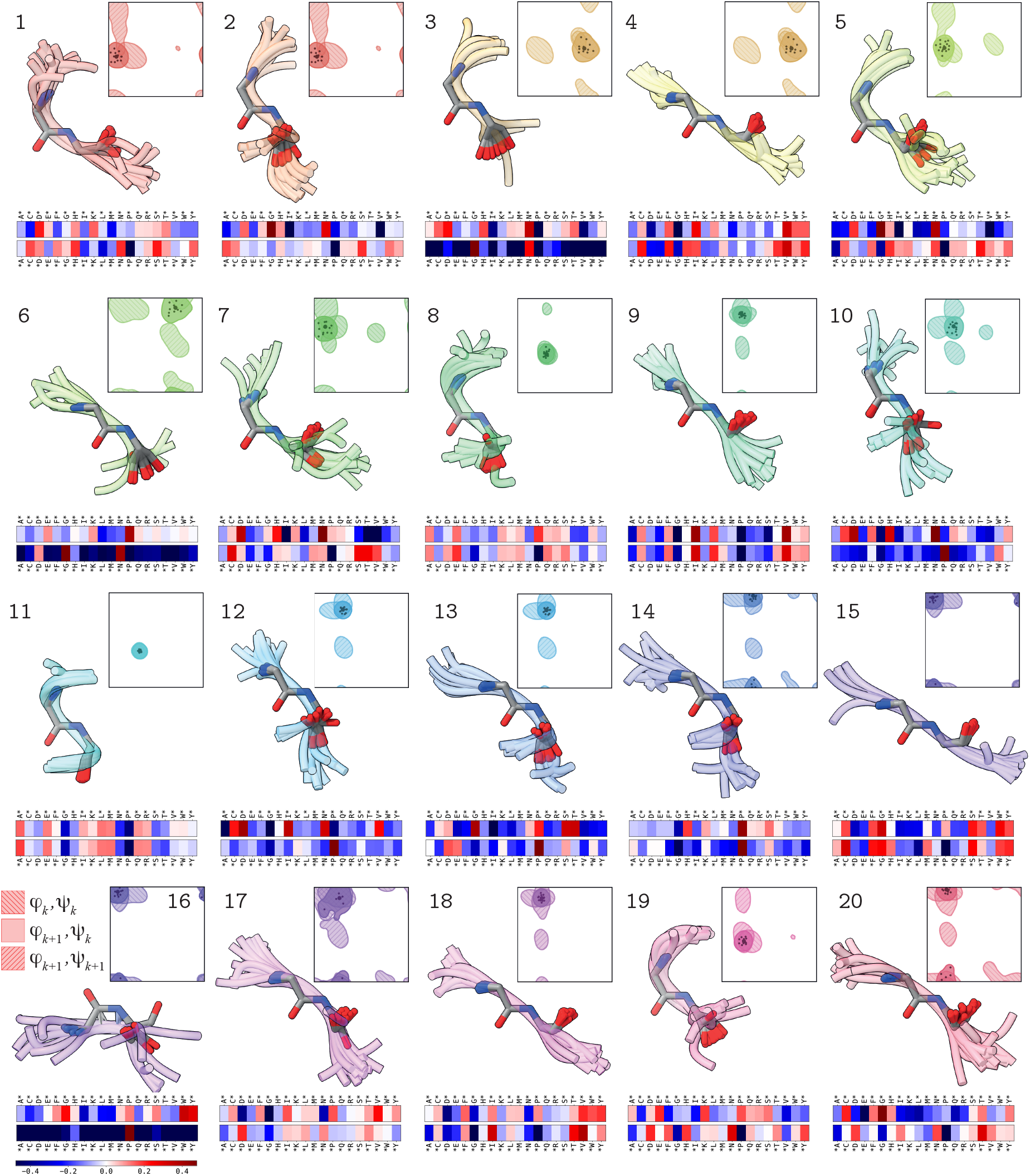
5,648,759 consecutive amino acid pairs from 24,875 distinct PDB protein chains with resolution above 1.8Å were brought into canonical alignment (Fig. S1) and clustered into 20 clusters using the *k*-means algorithm. Each numbered panel shows the backbones of 10 representative structures within the correspondingly numbered cluster, in the context of surrounding two amino acids (one on each side of the pair). The inserts show the regions occupied by each cluster in the three marginal plots (solid fill indicates the proposed cross-bond Ramachandran plot, while hatched fills depict the “conventional” plots). The colored bars below each cluster indicate the log propensity of different amino acid pairs of the form `X*` and `*X` to belong to the cluster.

**Fig. S3.**
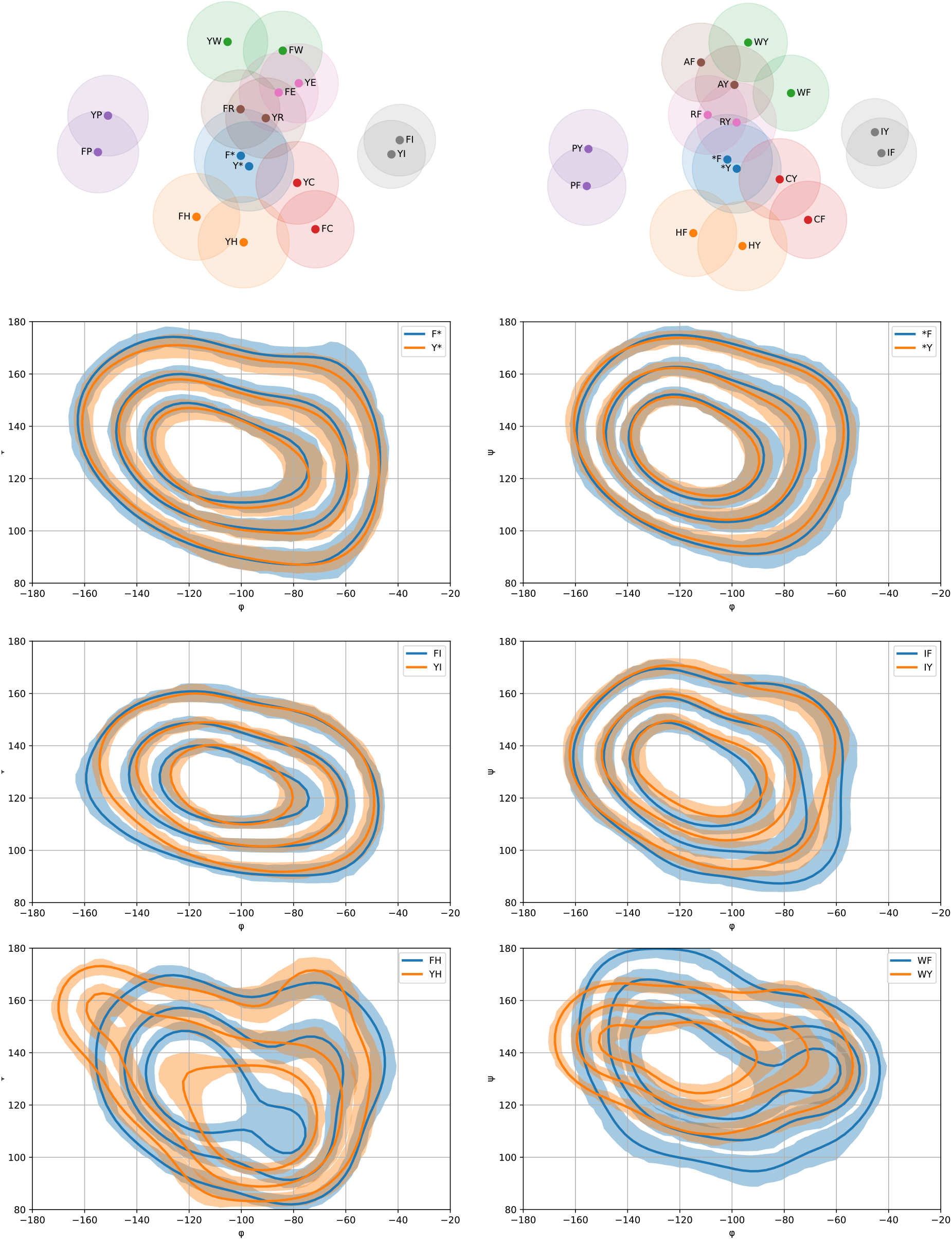
The impact of left and right amino acid context on the proposed cross-bond angle distributions. *Top*: L_1_ distances between the cross-bond distributions of pairs of amino acids of the form `FX` and `YX` (left), and `XF` and `XY` (right); `X=*` stands for no conditioning on the left/right context. Each point represents an amino acid pair, Euclidean distances between points correspond to L_1_ distances between the (ψ_*k*_, φ_*k*+1_) distribution of each pair, and shaded circles indicate 90% confidence regions. Note that `F*` and `Y*` as well as `*F` and `*Y` have nearly identical distributions, which is unsurprising as phenylalanine and tyrosine are chemically and structurally similar amino acids, and `F<>Y` is considered a conservative substitution. Some contexts like proline make the distribution substantially different (observe the location of the `YP, FP` and `PY, PF` pairs compared to `Y*, F*` and `*Y, *F`, respectively). More interestingly, some contexts like histidine on the right and tryptophan on the left make the distributions of the corresponding YH, FH and WY, WF pairs substantially different. *Bottom:* density plots of the (ψ_*k*_, ϕ_*k*+1_) dihedral angles of conservatively substituted `F<>Y` in different left and right contexts. Contours containing 25%, 50%, and 75% probability are shown; shaded regions indicate 90% confidence. Note the substantial difference between `YH, FH` and `WY, WF` pairs.

**Fig. S4.**
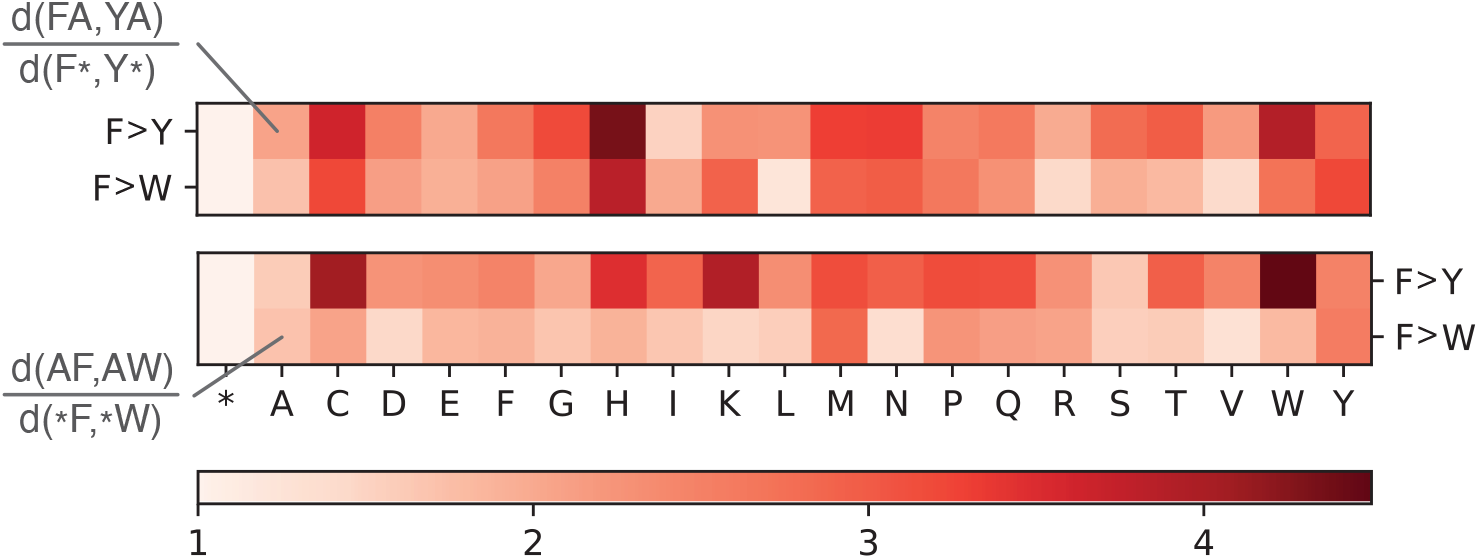
Normalized distribution distances for `F>Y` and `F>W` conservative substitutions in different right (top row) and left (bottom row) contexts. Normalized distance between a pair `FX` and `YX` is calculated as the ratio between d(`FX, YX`) and d(`F*, Y*`), where d stands for the L1 distance between (ψ_*k*_, ϕ_*k*+1_) dihedral angle distributions, and `F*, Y*` refer to `F` and `Y` not conditioned by any right context.

**Fig. S5.**
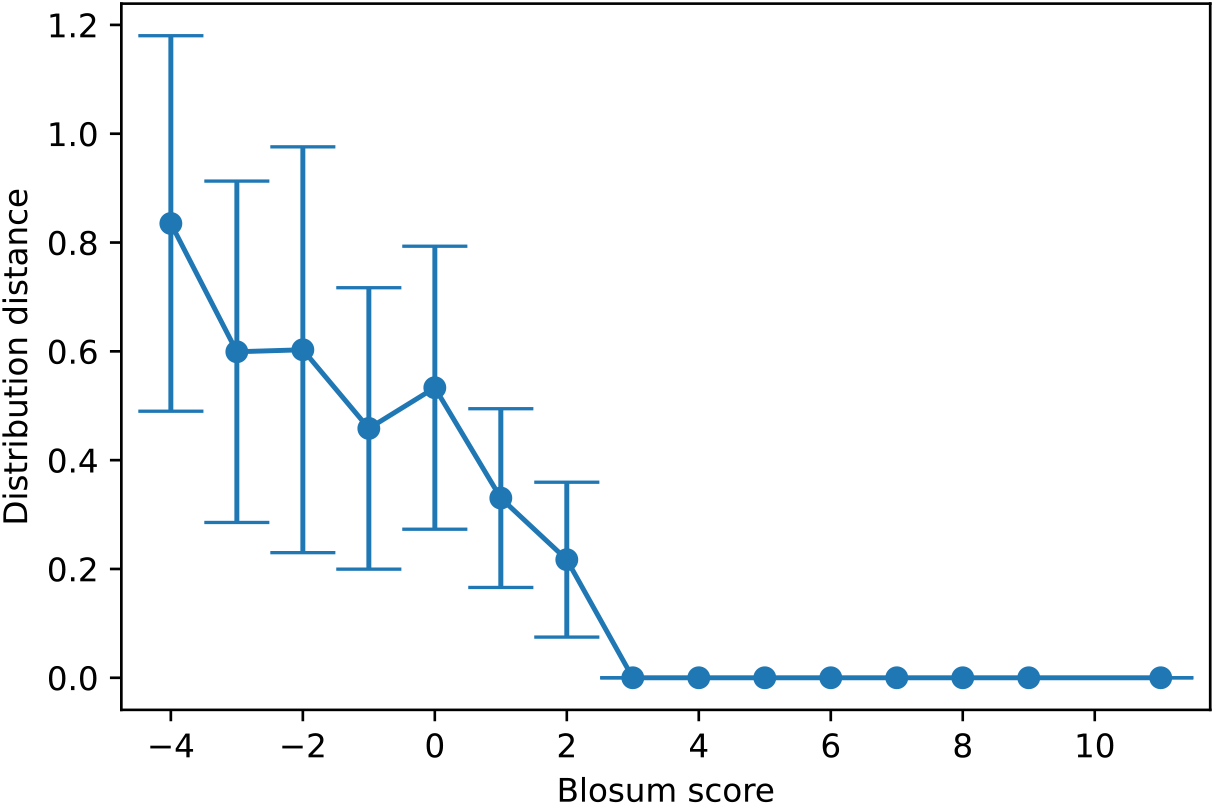
Dihedral angle distribution distances and BLOSUM62 scores are correlated. The distance between dihedral angle (ϕ_*k*_, ψ_*k*_) distributions of single amino acids is predictive of their mutation distance as embodied by the BLOSUM62 score.

**Table S1.**
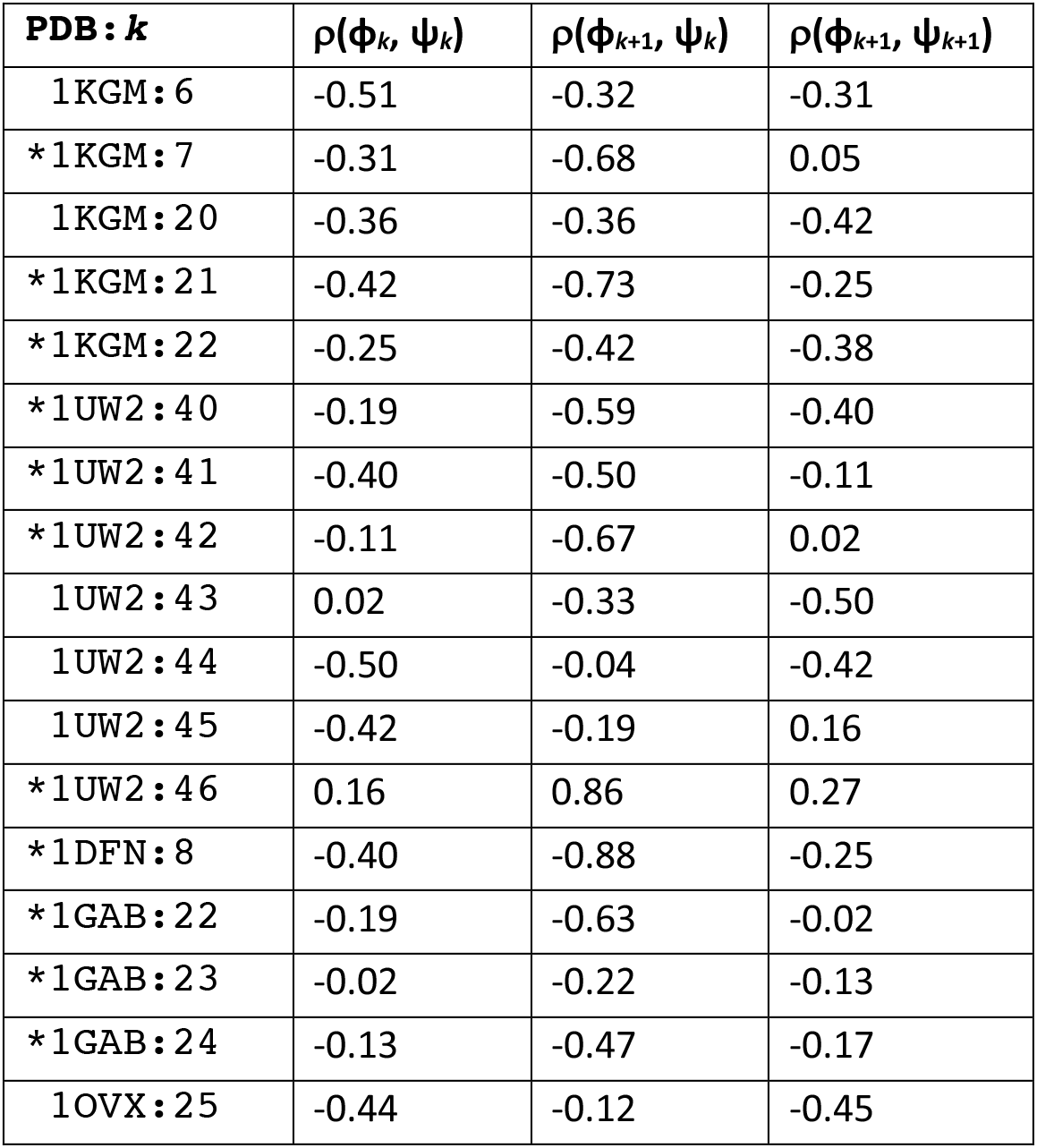
The cross-bond dihedral angle pair trajectories are at least as correlated as the conventional dihedral angle pair. Correlation coefficients ρ between different pairs of dihedral angles in molecular dynamics simulations of different non-interacting amino acid pairs within protein structures. *indicates examples for which the absolute value of the correlation coefficient is greater for the cross-bond dihedral angle pair.

